# Using in vivo calcium imaging to examine joint neuron spontaneous activity and home cage analysis to monitor activity changes in mouse models of arthritis

**DOI:** 10.1101/2024.08.28.610097

**Authors:** George L Goodwin, Alina-Cristina Marin, Julia Vlachaki Walker, Carl Hobbs, Franziska Denk

**Affiliations:** Wolfson Sensory, Pain and Regeneration Centre (SPaRC), King’s College London, SE1 1UL, UK

**Keywords:** Pain, Spontaneous activity, *In vivo* calcium imaging, Home cage analysis, Antigen induced arthritis, Partial medial meniscectomy

## Abstract

**Background:** Studying pain in rodent models of arthritis is challenging. For example, assessing functional changes in joints neurons is challenging due to their relative scarcity amongst all sensory neurons. Additionally, studying pain behaviors in rodent models of arthritis poses its own set of difficulties. Commonly used tests, such as static weight-bearing, often require restraint, which can induce stress and consequently alter nociception. The aim of this study was to evaluate two emerging techniques for investigating joint pain in mouse models of rheumatoid and osteoarthritis: *In vivo* calcium imaging to monitor joint afferent activity and group-housed home cage monitoring to assess pain-like behaviors. Specifically, we examined whether there was increased spontaneous activity in joint afferents and reduced locomotor activity following induction of arthritis.

**Methods:** Antigen induced arthritis (AIA) was used to model rheumatoid arthritis and partial medial meniscectomy (PMX) was used to model osteoarthritis. Group-housed home cage monitoring was used to assess locomotor behavior in all mice, and weight bearing was assessed in PMX mice. *In vivo* calcium imaging with GCaMP6s was used to monitor spontaneous activity in L4 ganglion joint neurons retrogradely labelled with fast blue 2 days following AIA and 13-15 weeks following PMX model induction. Cartilage degradation was assessed in knee joint sections stained with Safranin O and fast green in PMX mice.

**Results:** Antigen induced arthritis produced knee joint swelling and PMX caused degeneration of articular cartilage in the knee. In the first 46 hours following AIA, mice travelled less distance and were less mobile compared to their control cage mates. In contrast, no such differences were found between PMX and sham mice when measured between 4-12 weeks post-surgery. A larger fraction of joint neurons showed spontaneous activity in AIA but not PMX mice. Spontaneous activity was mostly displayed by medium-sized neurons in AIA mice and was not correlated with any of the home cage behaviors.

**Conclusion:** Group-housed home cage monitoring revealed locomotor changes in AIA mice, but not PMX mice (with n=10/group). *In vivo* calcium imaging can be used to assess activity in multiple retrogradely labelled joint afferents and revealed increased spontaneous activity in AIA but not PMX mice.

## Introduction

Current analgesic treatments for arthritis pain are not sufficient. For instance, first line drugs for osteoarthritis (OA) pain are either ineffective and/or poorly tolerated (1). Thus, we urgently need to better understand the mechanisms that lead to the development of pain in order to develop better analgesic medications. One limitation of this development is the comparatively low number of studies examining pain in preclinical models of arthritis. For instance, <10% of papers studying OA in mice examined pain between 2018-2023, and very few studies examined functional changes in neurons (2). This omission may appear strange, given that pain has a huge impact on individuals living with arthritis (3); this is until one is reminded of how challenging it is to study joint pain in rodents.

Functional changes to sensory neurons are particularly difficult to assess and traditionally have had to rely on highly specialized electrophysiological techniques (4–6). These are very low-throughput and, in the context of arthritis, have to obtain recordings from the very small proportion of neurons that innervate joints e.g. only ∼3-5% of lumbar sensory neurons project to the intra-articular space of the mouse knee (7). *In vivo* calcium imaging is a relatively new approach for assessing sensory neuron function, which we and others have utilized to study models of arthritis (8–10). However, thus far, only global changes in all types of sensory neurons within a given ganglion have been examined; important functional changes in joint afferents specifically may therefore have been ‘diluted’, seeing as this population is relatively small.

Another issue with studying pain in pre-clinical models of arthritis is the relative scarcity of robust and reproducible behavioral assays. Mice are now more commonly used than rats in pain behavioral studies (11). However, they are a more challenging species in this regard, because they require longer habituation times and are more prone to experimenter induced stress compared to rats (11, 12). Static weight bearing behavior is purported to be the ‘gold-standard’ for assessing joint pain in mice (13, 14). This test normally involves restraint, which can lead to stress-induced alterations of behavioral responses, such as modulation of nociception (15). This might explain why the reproducibility of static weight bearing in surgical models of OA in mice is mixed: some groups find significant shifts in weight bearing at late timepoints (14, 16), whereas others do not (17, 18). Larger weight bearing shifts are seen in arthritis models that induce substantial knee swelling, like AIA (18), however, these experiments are difficult to blind due to the visibly obvious phenotype induced by the model.

Here, we set out to establish whether *In vivo* calcium imaging and group-housed home cage analysis could be used to overcome some of the challenges of studying joint pain in mice. Specifically, our objectives were to: 1) assess whether retrograde labeling and *In vivo* imaging could be used to measure spontaneous activity in joint-specific neurons, and 2) evaluate if arthritis models induce changes in overall locomotor activity in the home cage. We hypothesized that an increased proportion of joint neurons would show spontaneous activity in the AIA and PMX models and that arthritic mice would exhibit reduced activity levels following model induction. The models were chosen to reflect the two conditions most likely to cause joint pain in people (osteo- and inflammatory arthritis) and to assess the sensitivity of our methods; thus, in the AIA model, joint swelling and consequent functional and behavioral changes are very pronounced at acute timepoints (19, 20); in contrast, the PMX model has a more subtle behavioral phenotype, and functional changes in sensory neurons have not been studied (14, 17).

Using the AIA model of rheumatoid arthritis and the PMX model of osteoarthritis in mice, we can record from multiple back-labelled joint afferents simultaneously. We observed increased spontaneous activity in AIA but not PMX mice. Home cage analysis showed significant reductions in activity following methylated bovine serum albumin (mBSA) injection in AIA mice, but not following PMX surgery.

## Materials and Methods

### Animals

Adult C57BL/6J male and female mice (n =69; Charles River, UK) were used in this study. Mice were housed on a 12/12 h light/dark cycle with a maximum of 4 mice per cage, with food and water available *ad libitum*. All experiments were performed in accordance with the United Kingdom Home Office Animals (Scientific Procedures) Act (1986). Animals undergoing behavioral experiments were housed in cages of 4 and were randomly assigned to treatment groups with 2 treatment and 2 controls in each cage. The experimenter assessing behavior was blinded to the treatment group during testing and analysis.

### Administration of calcium indicator

We utilized the genetically encoded calcium indicator GCaMP6s for imaging sensory neuron activity (21). GCaMP6s was delivered to sensory neurons via an adeno-associated viral vector of serotype 9 (AAV9), which was administered to mouse pups at post-natal day P2-P5 as previously described (22). Briefly, groups of 3-4 pups were separated from their mother and 6 µl of AAV9.CAG.GCaMP6s.WPRE.SV40 (Addgene, USA) was injected subcutaneously in the nape of the neck, using a 10 µL Hamilton syringe with a 30G needle. The pups were then returned to their mother until weaning at P21, when they were separated and housed with same sex litter mates until being used for *In vivo* imaging from 12 weeks after the injection.

### Retrograde labelling of knee joint afferents

10-12 week old mice were anesthetized using ketamine and xylazine (75mg/kg and 1mg/kg) and placed on a heating blanket. After shaving the knee and wiping it with 70% ethanol, the patellar tendon was identified and marked by running a 27g needle in the gap beneath the patella. A Hamilton syringe with a 30g needle attached was inserted through the patella tendon at a depth of 2-2.5 mm to deliver the retrograde tracer fast blue (FB; 2 μl 2% in H2O; Polysciences) into the articular space. Mice were then left on a heating blanket for 1.5 hours, while still under anesthesia, to ensure that fast blue was up taken into joint afferents. After reversal of anesthesia with antisedan (1mg/kg), mice were allowed to recover for 1.5 hours in a heated chamber before being returned to their home cage.

### Partial medial meniscectomy model

Twelve week-old male C57BL6/J mice (Charles River, UK) were randomized to undergo surgical destabilization by partial meniscectomy or sham surgery as previously described (23). Briefly, animals were placed under general anesthesia by inhalation of isoflurane (Vetpharma, Leeds, UK); 3% induction, 2% maintenance in 1-1.5 L/min O2. 0.1 mg/kg buprenorphine (Vetergesic Alstoe Animal Health, UK) and 1mg/kg butorphanol (Zoetis,UK) were administered subcutaneously for peri- and post-operative analgesia, respectively. For PMX surgery, the knee joint was opened and the meniscotibial ligament was transected and approximately 1 mm of the medial meniscus removed. For the sham operation, the knee joint was opened to expose the meniscotibial ligament, then closed with sutures.

### Antigen induced arthritis model

Mice were immunized using an emulsion of CFA (3.3 mg/ml; Sigma, UK) and mBSA (40 mg/ml; Sigma, UK), as described previously (24). Briefly, mice were anesthetized using 2 % isoflurane, and 100 µl of the emulsion was subcutaneously injected at the base of the tail and in the right flank (50 µl each). Mice were then allowed to recover and were returned to their home cages. Seven days after immunization, mice were anesthetized with isoflurane, and 2.5 µl of mBSA (200µg) was injected into the left knee joint using a Hamilton syringe with a 30G needle. Two days post-injection, the size across ipsilateral and contralateral knee joints was measured using a digital caliper.

### Static weight bearing

Weight bearing behavior was assessed using a Linton incapacitance tester. Mice were acclimatized to the experimenter, room and weight bearing apparatus on at least 3 separate occasions prior to baseline testing. For each test session, the force exerted over a 1s period by each hind paw was measured five times. Measurements from each paw were averaged, and the percentage of weight born on each side was calculated. Weight bearing was assessed at baseline and at 2-week intervals after osteoarthritis surgery.

### Home cage analyzer

Mice were microchipped in the lower flank (identichip, Animalcare Ltd, UK) whilst anesthetized for the fast blue administration. Groups of four mice (n=2 treatment and n=2 control for RA and OA models and n=4 for naïve controls) that were housed in their home cages (GM500, Tecniplast) were placed into a home cage analyzer (HCA, Actual Analytics, UK) (25), and recorded at baseline and different time points after induction of the arthritis models. Time spent mobile, distance travelled, time spent isolated - defined as being over 20cm away from nearest mouse, and average distance separated from nearest cage mate were measured by the HCA system using the position of the animal in cage, as localized using RFID readers below the cage. Measurements were aggregated over 1h time bins for each animal. To compare behavior before and after model induction, percentage change from baseline was calculated for light (07:00–19:00) and dark phases (19:00–07:00) by averaging measurements over complete 12-hour periods at each timepoint and comparing them to baseline averages. Only full 12-hour periods (19:00–07:00 for dark, 07:00–19:00 for light) were used. Statistical analysis was performed on this percentage change data. We did not analyze the anxiety-related behaviors, such as thigmotaxis, because the GM500 home cage is too small and thus not sensitive enough to detect changes in these parameters (communication from Actual Analytics).

### *In vivo* imaging of sensory neuron activity using GCaMP6s

*In vivo* imaging was performed as previously described (9). Briefly, mice were anesthetized using a combination of drugs: 1-1.25 g/kg 12.5% w/v urethane administered intraperitoneally and 0.5-1.5 % isoflurane delivered via a nose cone. Body temperature was maintained close to 37°C. An incision was made in the skin on the back, and the muscle overlying the L3-L5 DRG was removed. Using fine-tipped rongeurs, the bone surrounding the L4 DRG was carefully removed. The exposure was then stabilized at the neighboring vertebrae using spinal clamps (Precision Systems and Instrumentation, VA, USA) attached to a custom-made stage. Finally, the DRG was covered with silicone elastomer (World Precision Instruments, Ltd) to maintain a physiological environment. Prior to imaging, we administered a subcutaneous injection of 0.25 ml 0.9 % sterile saline to keep the mouse hydrated. It was then placed under an Eclipse Ni-E FN upright confocal/multiphoton microscope (Nikon).

The ambient temperature during imaging was kept at 32°C throughout. All images were acquired using a 10× dry objective. A 488-nm Argon ion laser line was used to excite GCaMP6s, and the signal was collected at 500–550 nm. Time lapse recordings were taken with an in-plane resolution of 512 × 512 pixels and a partially (66%) open pinhole for confocal image acquisition. All recordings were acquired at 3.65 Hz. A Z-stack was taken at the end of the experiment using a 488-nm laser to excite GCaMP6s and a 405-nm laser to excite fast blue. Stacks were taken every 4 µm over a typical depth of 80µm with the pinhole closed and channel series mode on to ensure no bleed through of emission signal between channels.

### Imaging data analysis

The researcher analyzing the data was blinded to the treatment group. Timelapse recordings were concatenated and scaled to 8-bit in Fiji/ImageJ, Version 1.53. The image analysis pipeline Suite2P (v 0.9.2)(26) was utilized for motion correction, automatic region of interest (ROI) detection and signal extraction. Further analysis was undertaken with a combination of Microsoft Office Excel 2013, Matlab (2018a) and RStudio (Version 4.02). A region of background was selected, and its signal subtracted from each ROI. To generate normalized ΔF/F0 data, the ROMANO toolbox ProcessFluorescentTraces() function was utilized (27). This function uses the calculation: ΔF/F0 = (Ft-F0)/F0, where Ft is the fluorescence at time t and F0 is the fluorescence average over the first 3 minutes of recording. ΔF/F0 is expressed as a percentage.

The area of each neuron in the L4 DRG was estimated by using the radius extracted with Suite2P. Cells were grouped into small to medium (≤750µm^2^) and large-sized (>750µm^2^) neurons. Knee joint afferents (FB+ GCaMP6s+) were identified using a z-stack max projection of blue and green channels in ImageJ. The area of FB+ neurons was measured precisely using the free hand selection tool in ImageJ. Cells that could not clearly be identified and had substantial overlap with neighboring cells were excluded from the analysis.

A positive response to capsaicin was taken if the max signal during the 5-minute post-capsaicin period was 10 SD greater than the 2-minute baseline period. Spontaneous activity was assessed during the first 5 minutes of the recording using a trained machine learning classifier (9).

We also aimed to assess spontaneous activity across all L4 neurons, including those innervating the knee joint and other areas, such as the hind paw. To optimize neuron detection by the activity algorithm in Suite2p, we applied a pinch stimulus to the hind paw using serrated forceps, maximizing the number of neurons automatically identified.

### *In vivo* application of capsaicin and lidocaine

The sciatic nerve was exposed at the level of the midthigh prior to drug application. In some experiments, 50µl capsaicin (Sigma, UK; 1mM in 0.2% ethanol 0.9% saline) was applied to the nerve. Any residual capsaicin was then removed from the nerve and lidocaine hydrochloride (Sigma, UK; dissolved in 0.9 % w/v sodium chloride to a concentration of 74 mM) was applied to block activity.

### Knee joint histology and analysis

After transcardial perfusion with PBS, mouse knee joints were extracted and processed as previously described (23). Briefly, joints were fixed for 24 hours in 10% Formalin and decalcified for 10 days in 0.5M EDTA (pH = 7.4). Following dehydration in 70-90% ethanol, joints were embedded in paraffin wax and coronal sections were cut from the central compartment of the joint using a microtome (≥ three slides with 5 x 5µm sections; 66 µm slide intervals).

Sections were deparaffinized with xylene and hydrated with alcohol to water. Slides were stained with hematoxylin and eosin and then washed for 10 mins with tap water. After that, slides were stained with 0.05% fast green solution for 5 mins followed by a rinse with 1% acetic acid solution for 10–15 seconds. Finally, slides were stained in 0.1% Safranin O solution for 5 minutes and then washed briefly twice with distilled water, followed by two washes with 95% ethyl alcohol, and then immediately placed into a 60°C Oven. After 1 hour, slides were removed and then placed into a mounting Xylene pot for 5 mins before being cover-slipped using DPX mounting medium.

Sections were imaged in brightfield mode at 10x using a slide scanner (Axioscan 7, Zeiss). The best section from each slide, i.e. that without substantial folding/tearing, was selected for analysis. Three to six sections per joint were scored using the Osteoarthritis Research Society International (OARSI) system (28) by two researchers who were blinded to treatment group. The average score for medial and lateral compartments was calculated. In one joint sample from the sham group the lateral compartment was completely missing due to incorrect orientation of the sample.

### Quantification and Statistical analysis

Graphing and statistical analysis was undertaken with a combination of Excel 2013, R Studio (Version 4.02), GraphPad Prism (Version 10) and SPSS (Version 29.0). Details of statistical tests and sample sizes are recorded in the appropriate figure legend, while observed effects & sensitivity analyses are provided in Supplementary Table 1.

### Data loss and exclusion

See supplementary methods for information on data loss and exclusion.

## Results

### Partial medial meniscectomy causes loss of articular cartilage and antigen induced arthritis induces joint swelling

Histopathological analysis of the knee joints from PMX mice showed substantial loss and damage of articular cartilage compared to sham controls (F (1, 17) [Treatment] = 45.1, P<0.001; repeated measures 1-Way ANOVA; n=9-10/group; Figure 1A-D and Supplementary Figure 1). The mean OARSI score in PMX mice on the medial side (4.26 +/-0.37; Figure 1B&C) was significantly greater than the medial side of sham mice (0.91 +/-0.17; p= 0.0000004; Bonferroni post hoc comparison; Figure 1B&D). As expected, cartilage loss was mostly confined to the medial portion of the joint in PMX mice (F (1, 17) [Knee side*Treatment] = 32.0, P<0.001; repeated measures 1-Way ANOVA; Figure 1C): the mean OARSI score in the medial side (4.26 +/-0.37) was significantly greater than the lateral side (1.69 +/-0.26; p= 0.000001; Bonferroni post hoc comparison).

**Figure 1.**
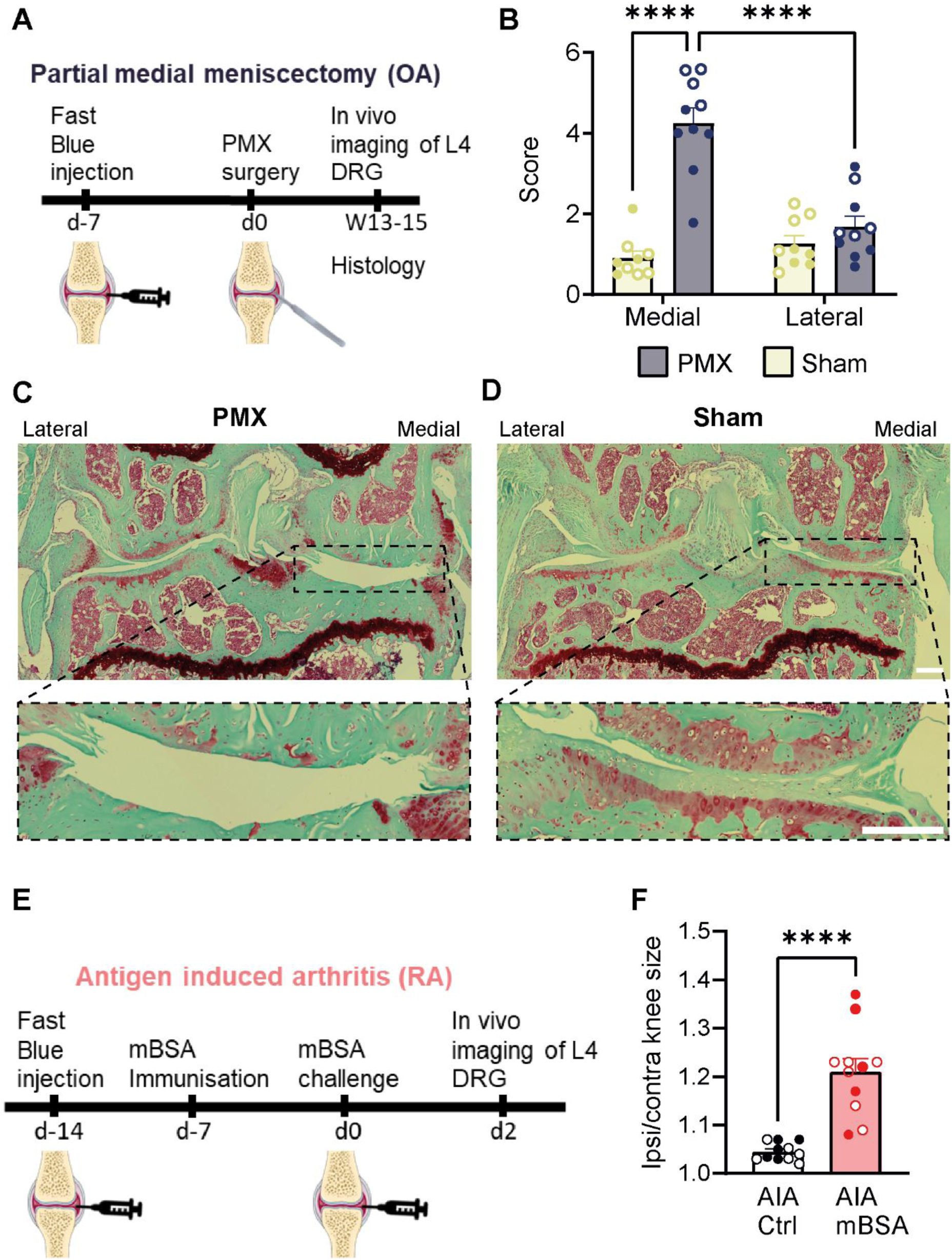
Partial medial meniscectomy causes loss of articular cartilage and mBSA injection causes joint swelling. **Legend for** Figure 1. A) Schematic showing experimental time course for the partial medial meniscectomy model of osteoarthritis. B) The average OARSI score to measure cartilage degradation for the medial and lateral compartments of the knee joint. Note that a minimum OARSI score of 0.5 was given to all samples because there was a lack of red staining in the superficial cartilage in all sections. C&D) Example images of Safranin O & fast green-stained knee joint sections in PMX (C) and Sham (D) mice at 13-15 weeks post-surgery. Scale bar = 200µm. E) Schematic showing experimental time course for the antigen induced arthritis model of rheumatoid arthritis. F) Ratio of the ipsilateral to contralateral knee size in AIA mice. Each point represents data from an individual animal. PMX: n= 9-10 mice/group AIA: n=11 mice/group. males = closed circles, females = open circles. PMX data: *** p<0.001 **** p<0.0001, repeated measures ANOVA. AIA data: p<0.0001, unpaired t-test.

Both PMX and sham mice showed reduced red staining in the superficial cartilage, which may indicate proteoglycan loss (28). As this was unexpected in sham mice based on previous histopathological findings (28) and was not observed in naïve mice, we wondered whether the prior intraarticular injection of fast blue into the knee joint caused these changes. To test this idea, we injected fast blue into joints of naïve mice and found that there was indeed discoloration of the superficial cartilage 7 days following fast blue but not saline injection (Figure 2 A&B): the blue and green intensity ratios in the superficial cartilage vs. deep layer was significantly increased in fast blue compared to saline-injected mice (F (1, 5) [Group] = 151.8, P=0.00006; repeated measures 1-Way ANOVA, n=3-4/group; Figure 2 C).

**Figure 2.**
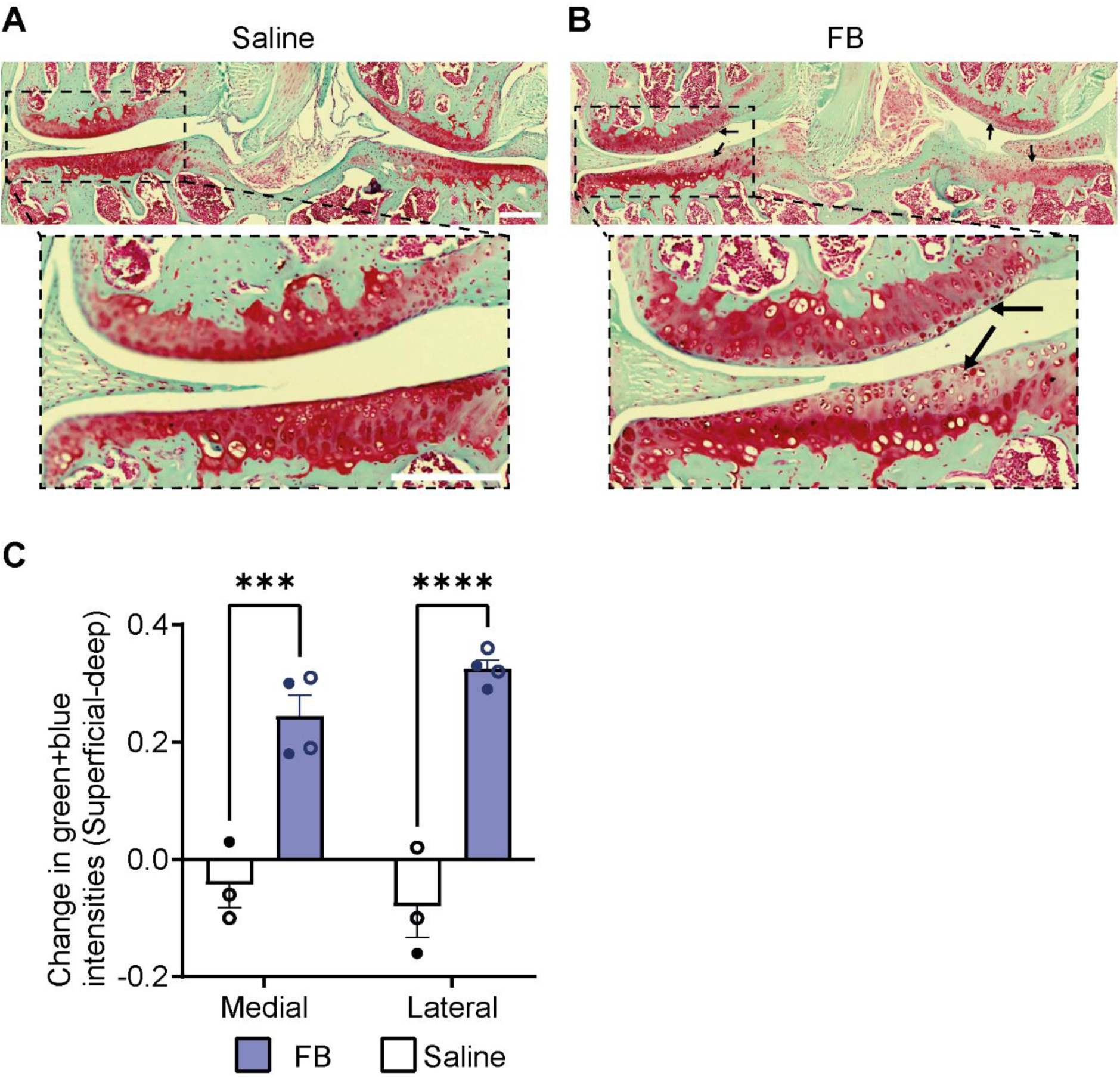
Safranin O staining of the superficial cartilage is altered following intraarticular injection of fast blue. **Legend for** Figure 2. A) and B) Example images of Safranin O and fast green stained knee joint sections following intraarticular injection of saline (A) and fast blue (B). The Black arrows show where there are changes to the intensity of the Safranin O staining in fast blue injected mice. Scale bar = 100µm. Note that the contrast has been altered in these images to highlight changes to the intensity of the Safranin O staining. C) Graph showing the difference in the blue and green: red intensity ratio between superficial and deep areas of the cartilage for medial and lateral compartments of the joint. A positive shift in the ratio indicates a higher proportion of blue and green staining compared to red in the superficial portion of the cartilage. Each point represents data from an individual animal. n=3-4 mice/group. males = closed circles, females = open circles. *** p<0.001 **** p<0.0001, repeated measures ANOVA.

In mice that were previously immunized with mBSA, 2.5 µl mBSA (200µg) injection into the knee joint caused a significant increase in swelling in the ipsilateral knee (Ipsilateral/contralateral size ratio = 1.21 +/-0.03) compared to mice injected with 2.5 µl saline (Ipsilateral/contralateral = 1.04 +/-0.01; p = 0.00008, n=11/group; unpaired t-test; Figure 1 E&F).

### Home cage analysis shows activity is reduced in antigen induced arthritis but not partial medial meniscectomy mice

We assessed whether group-housed home cage monitoring could be used to detect changes in activity following arthritis model induction (Figure 3A). As expected, mice were mostly quiescent during the light phase (07:00-19:00) and were more active during the dark phase (19:00-07:00; Figure 3B&D and Supplementary figure 2A-B). During the light phase, no differences were found in overall locomotor activity between AIA mBSA and control mice (F (1, 17) [Treatment group] = 1.6, P=0.23; repeated measures 1-Way ANOVA; n=9-10/group). However, during the dark phase there was a main effect of treatment (F (1, 17) [Treatment group] = 10.9, P=0.0001; repeated measures 1-Way ANOVA; n=9-10/group; Figure 3B-E). Specifically, mBSA injected mice travelled less distance (between subject effect F (1, 17) [Distance] = 25.8, p=0.00009; Figure 3 B&C) and were less mobile (between subject effect F (1, 17) [Mobility] = 25.3, p=0.0001; Figure 3 D&E) compared to saline injected mice. Following mBSA injection, there was a reduction in distance (-44.0%, +/-5.3) and mobility (-21.6%, +/-5.0) during the second dark phase, which was significantly different to the % change in saline injected mice at the same timepoint (distance = -22.2%, +/- 5.5; mobility = -0.7%, +/- 5.3; both p=0.011 Bonferroni post hoc comparison; Figure 3 C&E). In contrast, there were no significant changes to the amount of time spent isolated (between subject effect F (1, 17) [Isolation] = 2.4, p=0.14) and the average distance separated (between subject effect F (1, 17) [Separation] = 6.5, p=0.21; Supplementary figure 2 C-H).

**Figure 3.**
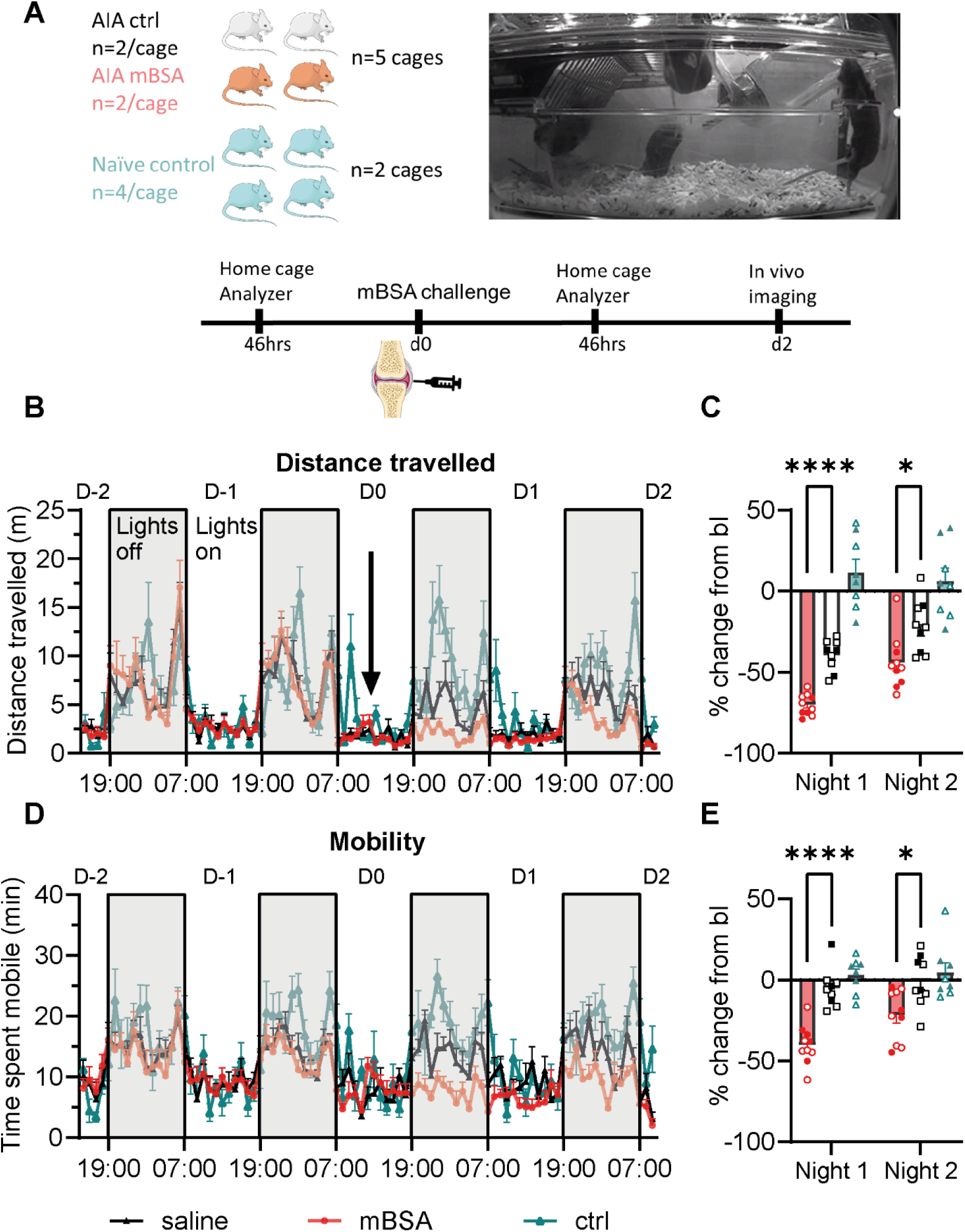
Home cage analysis shows that antigen induced arthritis mice are less active following mBSA injection. **Legend for Figure 3.** A) Schematic showing time course for the home cage analyzer experiments in AIA mice. Home cage analysis was performed in cages that contained 4 mice with n=2 mBSA and n=2 saline mice per cage, whereas naïve ctrl mice were assessed in groups of n=4. B-D) Graphs showing the total distance travelled (B&C) and time spent mobile (D&E) before and after intraarticular injections (indicated by arrow in B). Data in B) and D) are presented as mean +/- SEM in 1hr intervals. Data in C) and E) show the percentage change from baseline for the dark phase (19:00-07:00). Grey bars represent the dark phase. Home cage analysis data were collected from n = 10 mBSA and n = 9 saline mice, housed in n = 5 cages, grouped as described in A) and n = 8 naïve control mice in n= 2 cages. Males = closed circles, females = open circles. ****p<0.0001 *p<0.05, repeated measures 1-way ANOVA with Bonferroni adjusted post hoc comparisons.

There was no significant effect of treatment in PMX OA experiments during the light (F (1, 18) [Treatment group] = 18.9, P=0.12; repeated measures 1-Way ANOVA; n=10 group) or dark phases (F (1, 18) [Treatment group] = 0.5, P=0.72; repeated measures 1-Way ANOVA; n=10 group; Figure 4A-E), suggesting that changes to activity are not as severely impacted in this model. At 12-weeks following PMX surgery, the % change in distance (-12.8%, +/- 8.2) and mobility (-21.2 %, +/- 5.3) during the dark phase was similar to that of the sham group (distance = -16.4%, +/- 6.6; mobility = -17.8%, +/- 3.3; Figure 4 C&E). Similarly, the amount of time mice spent isolated and the average distance separated before and after surgery was comparable between PMX and Sham mice (Supplementary figure 3A-D).

**Figure 4.**
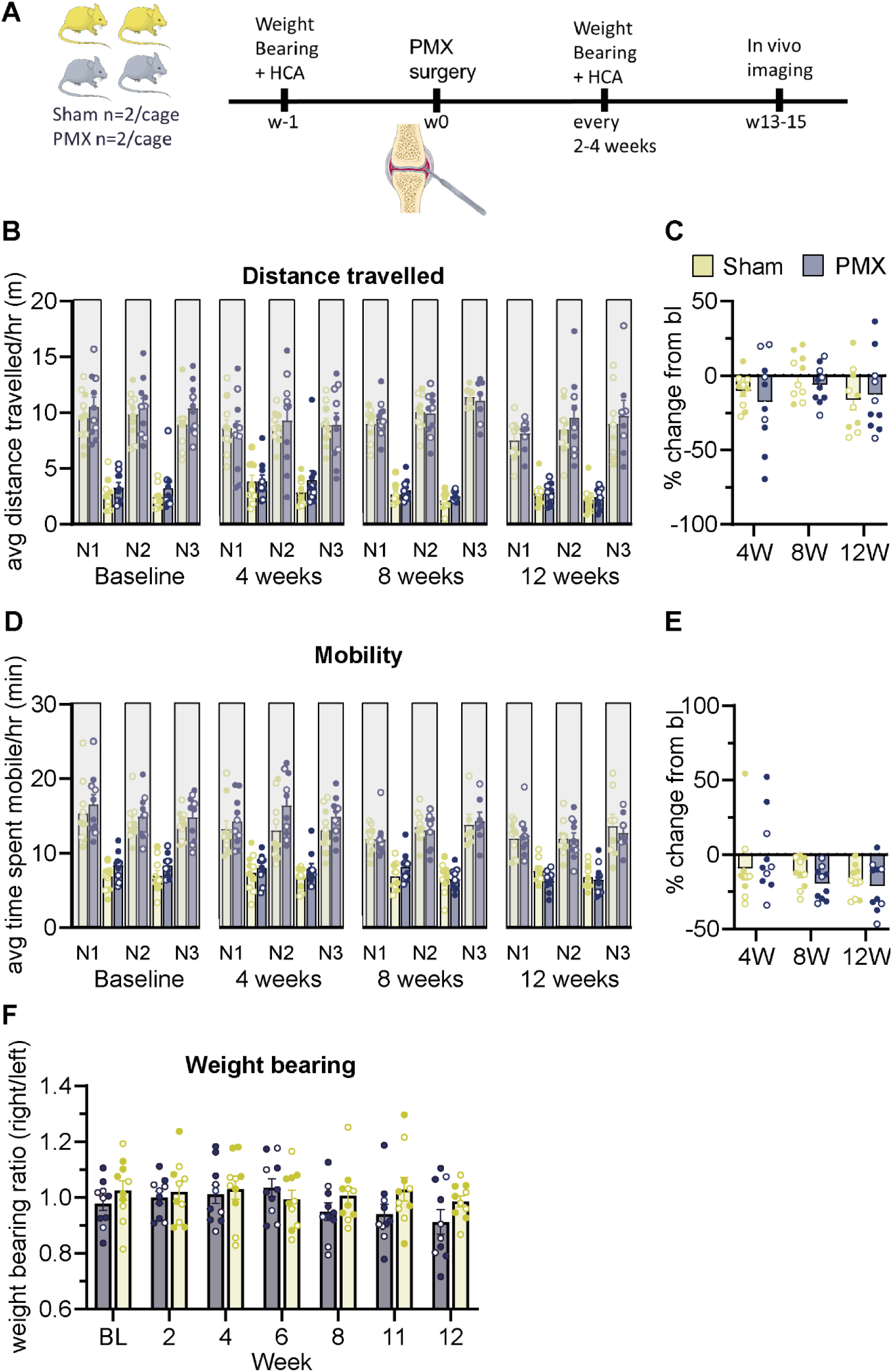
Home cage analysis shows that activity was similar in partial medial meniscectomy and sham mice. **Legend for Figure 4**. A) Schematic showing time course for the home cage analyzer experiments in OA mice. Home cage analysis was performed in cages that contained 4 mice with n=2 PMX and n=2 sham/cage. B-D) Graphs showing the total distance travelled (B&C) and time spent mobile (D&E) at baseline and following surgery. Data in B) and D) are presented as mean +/- SEM in 12hr intervals. Grey bars represent the dark phase. Data in C) and E) show the percentage change from baseline for the dark phase (19:00-07:00). F) Graph showing the weight bearing ratio (right/left hind limb) over time. Data are presented as mean +/- SEM. Home cage analysis assessed in n = 5 cages with n = 10 / group. Males = closed circles, females = open circles.

Weight bearing was also assessed in the PMX mice. PMX mice began to shift weight from their ipsilateral to contralateral hind limbs at 8 weeks following surgery (Ipsilateral/contralateral ratio mean = 0.95 +/- 0.03, Figure 4F). By 12 weeks, the ipsilateral/contralateral ratio in PMX and sham mice was 0.91 (+/- 0.04, n=10) and 0.99 (+/- 0.02, n=10), respectively (Figure 4F). However, significant differences were not found at any timepoint (F (6, 108) [Treatment x Time] = 1, P=0.46; repeated measures 1-Way ANOVA; n=10 group).

### *In vivo* imaging of retrogradely labelled joint afferents reveals increased spontaneous activity in mBSA injected mice but not PMX mice

We optimized our *In vivo* imaging approach for recording from fast blue retrogradely labelled joint afferents, i.e. those which innervate the synovium and ligaments, but not bone (29), and used it to assess spontaneous activity in AIA and PMX models (Figure 5A and Supplementary Figure 4). We focused on spontaneous activity because reports indicate that it is the likely neurophysiological correlate of unpredictable or ‘spontaneous’ pain (30, 31), which has a significant impact on quality of life in those with chronic arthritis pain (32).

**Figure 5.**
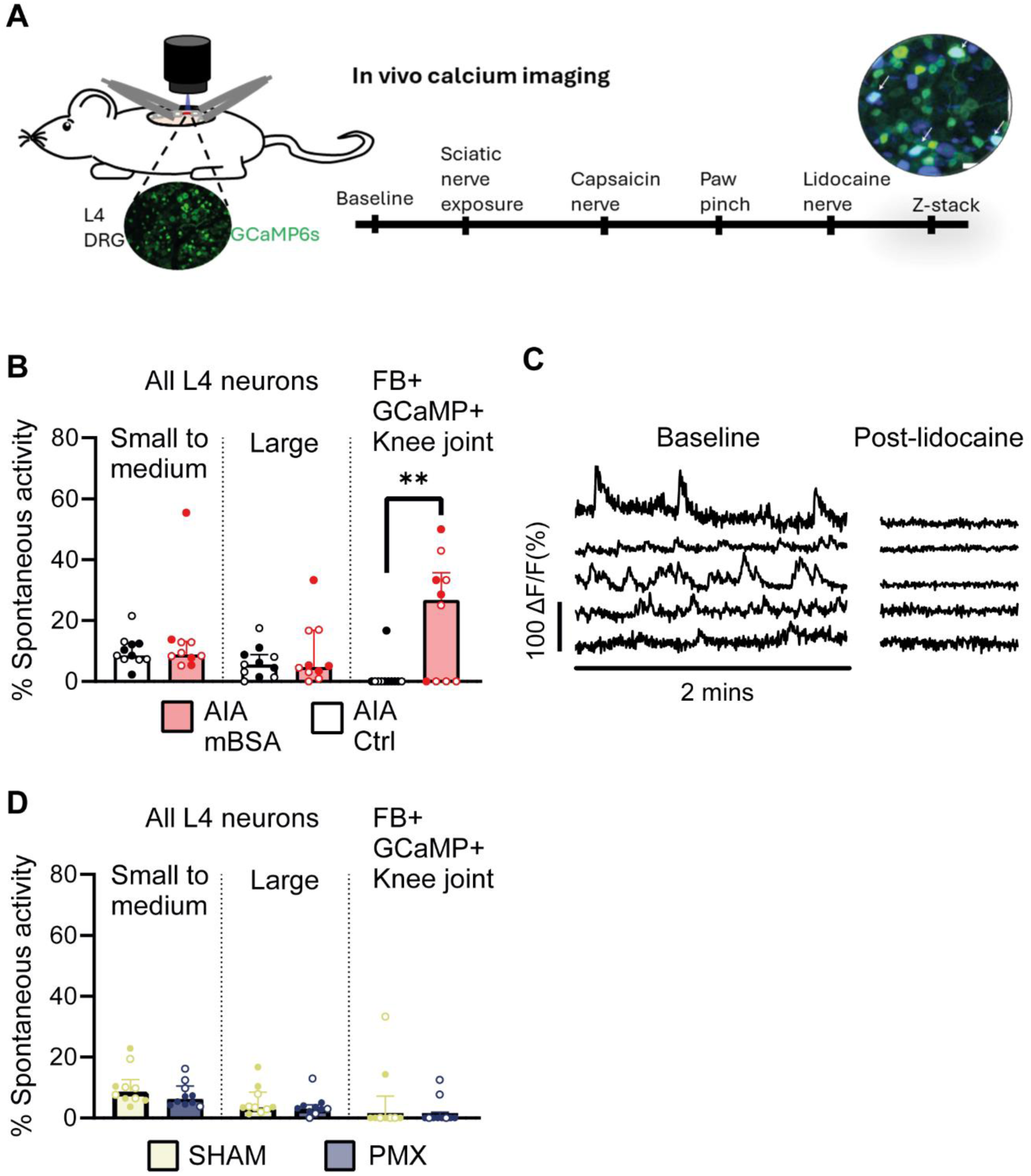
Increased spontaneous activity in joint afferents in mBSA injected mice but not in partial medial meniscectomy mice. A) Schematic showing *In vivo* calcium imaging experimental paradigm. B&D) Graphs showing the average proportion of neurons with spontaneous activity for small to medium, large and FB+ knee joints afferents in AIA (B) and PMX (D) experiments. C) Example calcium traces from spontaneously active neurons in mBSA injected AIA mice before (left) and after (right) lidocaine. Small to medium at <750µm² and large at ≥750 µm². Each point represents data from an individual animal. PMX weeks 13-15: n= 9-10 mice/group; AIA day 2: n=10-11 mice/group. males = closed circles, females = open circles. AIA data: ** p<0.01 Kruskal-Wallis test.

The proportion of double positive fast blue (FB+)/GCaMP6s+ cells among all GCamP6s labelled neurons was similar between AIA (1.7% +/-0.1) and PMX cohorts (2.1% +/- 0.2; Supplementary figure 4B). There was no spontaneous activity in FB+ joint neurons of naïve mice (0/15 neurons, n=3 mice) and only one active neuron in saline injected mice (1/19 neurons, median = 0%, n=3 mice), suggesting that the injection does not cause substantial abnormal firing in joint afferents. In AIA mice, mBSA injection caused a significant increase in the proportion of FB+ neurons with spontaneous activity (14/63 joint neurons, median = 26.8, IQR = 35.7), compared to saline injected controls (1/61 joint neurons, median = 0, IQR = 0; P = 0.009, Kruskal Wallis test; n=10-11/group; Figure 5B&C and Supplementary table 2). In contrast, the proportion of all L4 neurons with spontaneous activity was similar between mBSA and saline injected mice for small to medium- and large-sized neurons (Figure 5B and Supplementary table 2).

PMX was unlike AIA, with the levels of spontaneous activity in FB+ joint neurons at 13-15 weeks following PMX injury (2/75 neurons, median = 0 IQR = 1.9) similar to that of sham mice (2/55 neurons median = 0 IQR = 7.2; p =0.11, Kruskal Wallis test; n=10/group; Figure 5D and Supplementary table 2).

In some experiments, we aimed to determine if spontaneously active neurons express transient receptor potential cation channel subfamily V member 1 (TRPV1) by applying the agonist capsaicin (1mM) to the sciatic nerve. In TRPV1 expressing neurons, we anticipated that there would be depolarization i.e., increased calcium due to channel opening, followed by block (33, 34), which is largely thought to be dependent on TRPV1 activation (34, 35). However, some spontaneously active neurons were blocked without any prior depolarization (Supplementary figure 5 and supplementary table 3), suggesting that the concentration of capsaicin applied (1mM) may have caused block independent of TRPV1, such as through inhibition of voltage gated sodium channels (36). We were therefore unable to verify the proportion of spontaneously active joint neurons that express TRPV1.

### Spontaneously active neurons are predominantly medium-sized in mBSA injected-AIA mice

Retrogradely labelled joint neurons were mostly small to medium in size in AIA and PMX cohorts (Supplementary figure 6), as expected based on previous reports (29, 37). The only FB+ joint neuron with spontaneous activity in one of the saline-injected mice was small-sized (area = 237 µm^2^). Conversely, spontaneous activity in small-sized (<400 µm^2^) joint neurons was only recorded in one mBSA injected mouse out of a group of n = 6. In that animal, we recorded three small-sized spontaneously active FB+ neurons and unusually high overall levels of spontaneous activity (52.9% of all L4 neurons). Most other FB+ joint neurons with spontaneous activity following mBSA injection were medium-sized (9/14 neurons from n=6 mice; Figure 6 A&B, Supplementary video 1). The median area of spontaneously active neurons was 483 µm^2^ (IQR = 225). We under-sampled from the smallest sized neurons (<350 µm^2^) in AIA mBSA injected mice and therefore cannot rule out the possibility that there was also increased activity within this subgroup (Supplementary figure 6).

**Figure 6.**
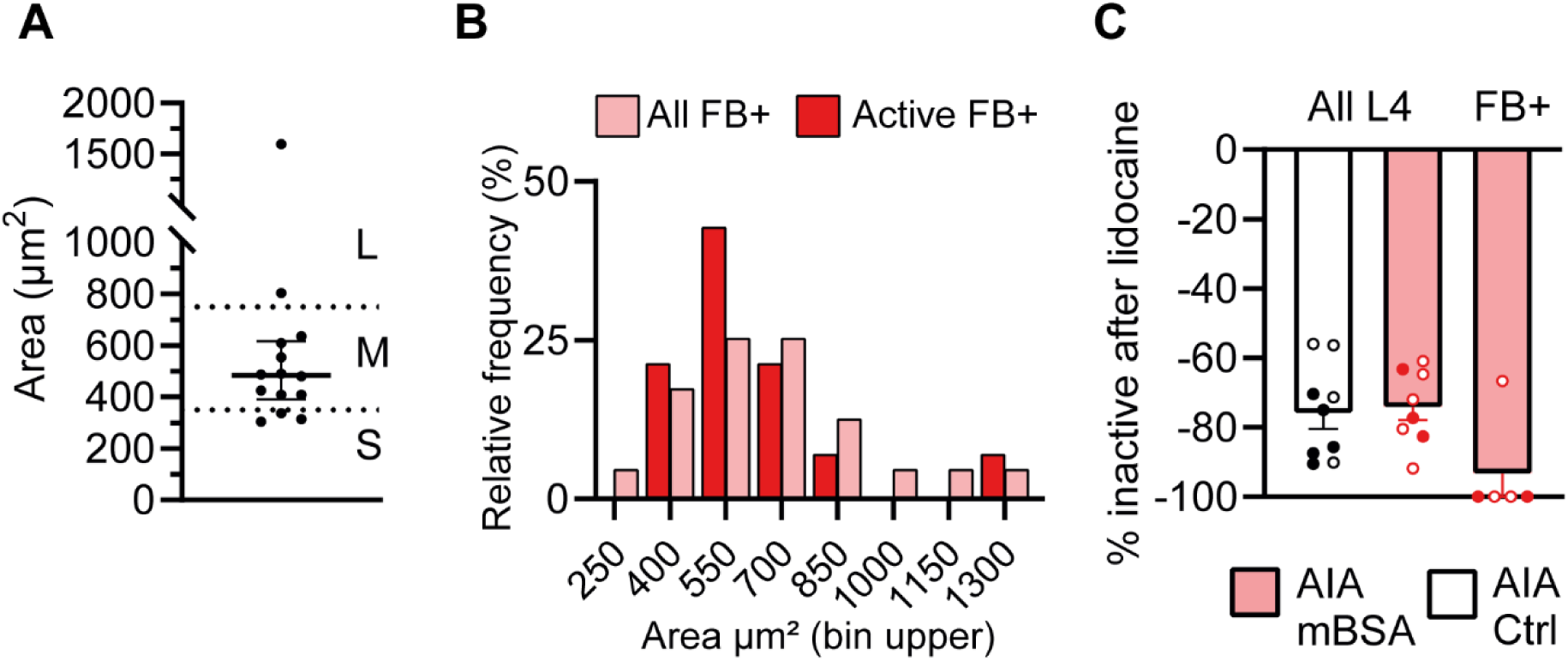
Most spontaneously active joint neurons are medium-sized and not active after peripheral lidocaine. A) Graph showing the size of the spontaneously active FB+ joint neurons in mBSA injected AIA mice. B) Size histogram showing the relative frequency of all FB+ joint neurons in mBSA injected mice (n=63) and those with spontaneous activity (n=14). Bin widths = 150 µm². The size of FB+ joint neurons was determined from z-stacks taken at the end of the recording. C) Graph showing the proportion of neurons that are not active following lidocaine application to the sciatic nerve.

Most spontaneously active neurons were silent after application of lidocaine to the sciatic nerve (n = 10/11 neurons; Figure 6C).

We also performed an exploratory analysis to determine if spontaneous activity in joint neurons was correlated with any of the home cage analyzer behavior parameters in AIA mBSA injected mice. No significant correlations could be detected in the n=8 AIA mice for which matched imaging and behavioral data were available (Figure 7). Of note, with this sample size, we are only well-powered to detect very large correlations (79% chance to detect Pearson’s r = 0.82+ at uncorrected p = 0.05).

**Figure 7.**
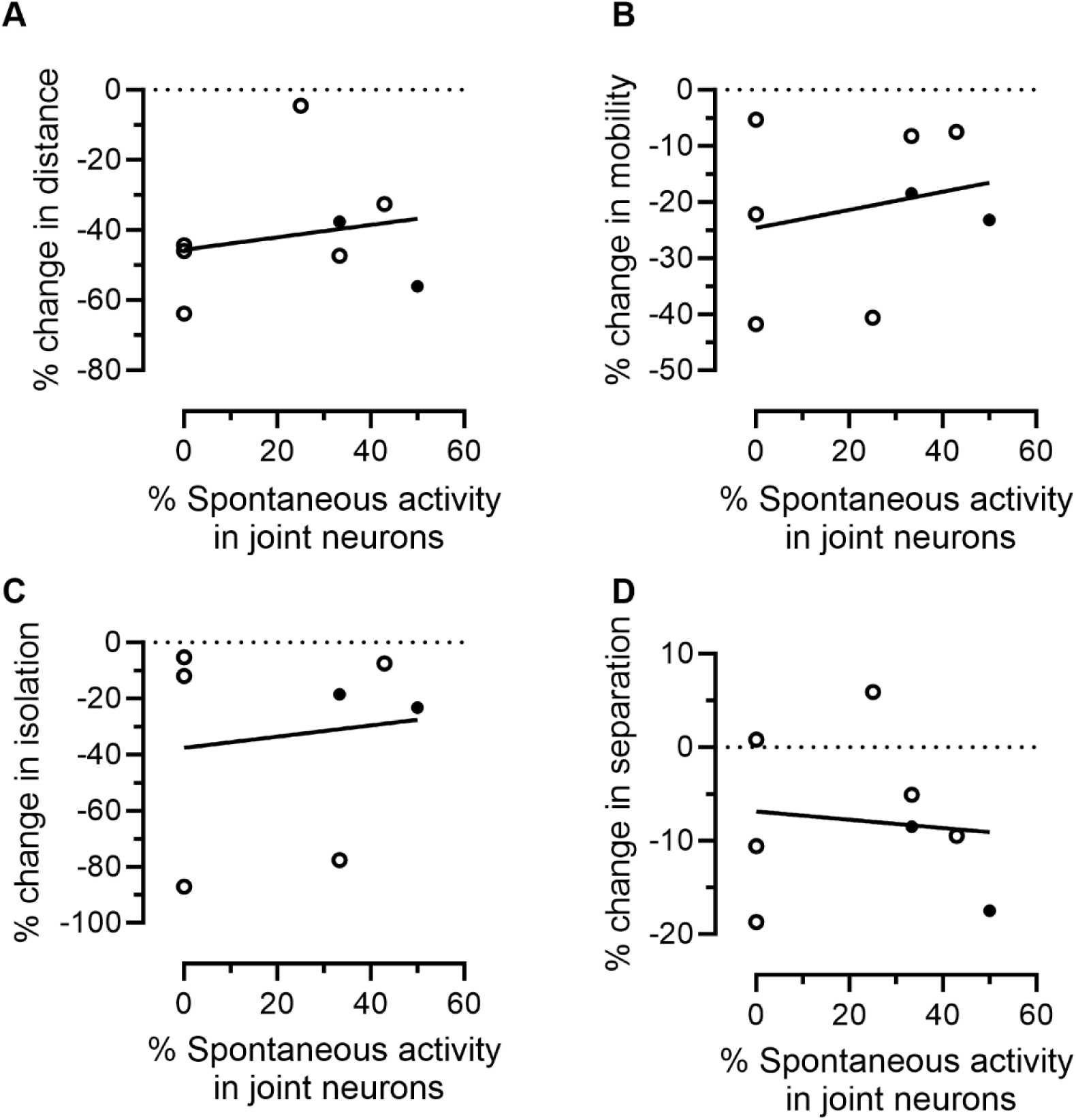
No correlations between joint neuron spontaneous activity and home cage analyzer behavior parameters in antigen induced arthritis mice. A-D) Graphs showing the relationship between the % of joint neurons with spontaneous activity and the % change in Distance (A), mobility (B), isolation (C), separation (D) on night 2 in mBSA injected AIA mice. The % change is between the baseline and night 2 data. N=8 mice, males = closed circles, females = open circles. Pearson’s correlations were as follows for distance (R^2^ =0.04, P=0.31), mobility (R^2^ =0.05, P=0.29), separation (R^2^ =0.01, P=0.4) and isolation (R^2^ =0.02, P=0.39). Data from n=8 mice.

## Discussion

In this study, we set out to determine if *In vivo* calcium imaging and group-housed home cage monitoring could be used for studying joint pain in models of RA and OA. We have shown that *In vivo* imaging can be used to monitor activity in multiple retrogradely labelled joint afferents (∼6/experiment) in the L4 DRG, which is greater than that sampled in a typical electrophysiology experiment (∼1-2/experiment (4)). In the AIA model of RA, we found increased spontaneous activity in joint afferents 2-days following intraarticular injection of mBSA, which is predominantly localized to medium-sized neurons. In contrast, spontaneous activity was not elevated in joint afferents at 13-15 weeks in the PMX model of OA. Group-housed home cage monitoring showed AIA mice were less active than their control cage mates in the first 46 hours following mBSA injection, whereas no differences were found between PMX and sham mice between 4-12 weeks post-surgery.

While previous studies have used *In vivo* imaging to assess neuronal activity globally in entire sensory ganglia in models of arthritis (8–10), our study is the first to monitor activity in retrogradely labelled FB+ joint afferents. These afferents are likely those that innervate the synovium and ligaments (38, 39), but not bone (29). We found increased levels of spontaneous activity in joint afferents in AIA mice, which is consistent with a previous electrophysiological study (19), and suggests that these afferents are involved in driving pain, in this model. We found that spontaneously active neurons were mostly medium-sized, which suggests they might be thinly myelinated Aδ-fibres. This appears in contrast to the C-fiber spontaneous activity reported by Qu et al.’s electrophysiological experiments (19). However, some medium-sized neurons are unmyelinated C-fibers (40), making it possible that our work agrees with their findings.

We studied the PMX model at a relatively ‘late’ timepoint because spontaneous pain, which is associated with increased spontaneous activity in neuropathic pain conditions (30, 31), is reported to become more severe in the later stages of OA in humans (32). It was therefore expected that spontaneous activity in joint afferents would be increased at a ‘late’ timepoint, i.e. week 13-15, in the PMX model. However, in contrast to AIA mice, intraarticular afferents did not have increased spontaneous activity at this ‘late’ timepoint following PMX. It is difficult to benchmark our data to past work, since there are very few other studies that have examined spontaneous activity in joint afferents in surgical models of OA. A recent study in rats found increased spontaneous activity in joint afferents at 4-weeks following medial meniscus transection (41). There are also reports of increased spontaneous activity in intraarticular afferents after monoiodoacetate injections (MIA) into the joint (4, 6); however MIA is qualitatively very different to surgical models of OA, with a much shorter time-course and a potential risk of direct neuronal damage (42).

Another known unknown is whether there might have been increased spontaneous activity in other afferent populations, such as bone, in late-stage PMX mice. The fast blue labelled neurons in our study are not likely to include bone afferents (29). Meanwhile, at whole DRG level, we would likely miss an increase in spontaneous activity if it was just restricted to bone afferents, as this fibre type constitutes a very small subset of all neurons, e.g. only 2% of L4 DRG innervate the tibia (43). This is experimentally supported by our AIA data, which show that differences in spontaneous activity in joint neurons (i.e. another rare sub-type of overall L4 DRG) are missed when all neurons are examined collectively (Figure 5). In addition to spontaneous activity, our *In vivo* imaging method has potential for examining changes to mechanical sensitivity of joint neurons. We are currently optimizing the best approaches for recording this, since it is well-established that joint neurons develop heightened sensitivity to mechanical stimulation in models of arthritis (5, 19, 41).

Beyond neuronal activity, we also examined mouse behavior using group-housed home cage monitoring. This removes many out-of-cage confounding factors, such as changes to environmental parameters and handling, allowing for a more unbiased assessment of behavioral changes in models of arthritis. Specifically, we assessed changes in overall locomotor activity, quantified as the total distance travelled and mobility during the 12h period when the mice are most active (dark phase). Our results indicate that this level of analysis is sensitive enough to detect changes in the AIA model of RA, but not in the PMX model of OA (with n=10 mice/group). AIA mice are presumed to be less active because they are in pain i.e. forces generated from behaviors that are normally regarded as innocuous, such as moving about, activate sensitized joint mechanical nociceptors. Mice may also have been inhibited in their movement by spontaneous joint neuron activity. While we found no correlation between this form of peripheral sensitization and home cage parameters, we were only powered to detect a very strong relationship between these variables. Our AIA data are consistent with previous studies using the LABORAS system in models of RA (14, 20), showing that overall locomotor activity is a reproducible measure of behavior change due to experimental RA. Meanwhile, our PMX home cage results mirror those of other studies using surgical models of OA in mice in the LABORAS system (14, 44), but contrast others (45, 46). Together, these data suggest that overall locomotor activity may not be a reproducible indicator of pain in surgical models of osteoarthritis, where the effects on behavior may be more subtle.

Are other parameters that can be assessed by home cage analyzers, such as climbing and rearing behavior, more sensitive for detecting changes to behaviors in surgical models of OA? Studies using the LABORAS system reported reduced climbing and rearing behavior in the destabilization of the medial meniscus (DMM) model (45, 46), while no significant differences were found examining the same parameters in the PMX model (14). However, as most pre-clinical arthritis studies are only powered to detect large effect sizes (with n≤10 mice/group), it is possible that the discrepancies between results are due to lack of power to detect more moderate effects.

Surprisingly, we also observed decreases in activity in our control mice for both models. It is not clear whether these changes were directly due to the control treatment i.e. the intraarticular injection or the sham surgery, or whether they were mediated through a social interaction effect i.e. the control mice move about less because their arthritic cage mates do.

Although we found a slight shift in weight bearing in PMX mice, it was not significantly different to the sham group, which is consistent with what others have reported (17, 18). The inconsistency of these results with the positive results that others have reported (14, 16) could stem from the restraint required for the test, which causes stress. A recent report found that less variable results can be produced when mice are unrestrained (47). Moreover, there could once again be a problem of statistical power: effects might not be large enough to be consistently detectable with the sample sizes commonly employed in the field.

Whilst the methods we employed have advantages over more traditional methods for assessing joint pain, there are limitations to our study. Although we are capable of sampling from ∼6 joint neurons/animal in our *In vivo* experiments, the number of neurons we record from is still relatively low, so results from our assays still need to be interpreted with some caution. Moreover, the terminal nature of DRG *In vivo* imaging only allows for the study of neurons at a single time point. It is therefore possible that increased levels of spontaneous activity might have been detected, if we were able to image mice repeatedly over the course of their arthritis. Whilst the number of mice used in our study (n=9-10) is similar to that of most other preclinical arthritis studies (14, 17), this sample size only provides the power to detect very large effects (e.g. with a parametric t-test, Cohen’s d in excess of 1.3). Thus, smaller effects may be present but will most probably be missed by the majority of current animal experiments in the field, including our own.

## Conclusion

We conclude that *In vivo* calcium imaging can be used together with retrograde labelling to assess functional changes in multiple joint neurons simultaneously. Using this technique, we found spontaneous activity in joint neurons was increased at an acute timepoint in AIA mice, but was not observed at a late timepoint in PMX mice. Group-housed home cage monitoring revealed changes to overall locomotor activity in AIA mice, showing that it can be used to detect arthritis-induced alterations to behavior. In contrast, no differences in overall locomotor activity were found in PMX mice, suggesting that this level of analysis may not be suitably sensitive to detect differences in behavior when using group sizes of n = 10.

## List of abbreviations

AIA: Antigen Induced Arthritis
CFA: Complete Freund’s Adjuvant
FB+: Fast blue positive
mBSA: Methylated Bovine Serum Albumin
OA: Osteoarthritis
OARSI: Osteoarthritis Research Society International
PMX: Partial Medial Meniscometry
RA: Rheumatoid Arthritis
TRPV1: transient receptor potential cation channel subfamily V member 1

## Declarations

### Ethics approval

All experiments were performed in accordance with the United Kingdom Home Office Animals (Scientific Procedures) Act (1986).

### Consent for publication

Not applicable.

### Availability of data and materials

Data have been shared via the SPARC portal under accession number https://doi.org/10.26275/xjgh-9rrj

### Competing interests

The authors declare that they have no competing interests.

### Funding

F.D. and G.G. are funded by Advanced Pain Discovery Platform UKRI MRC grants (MR/W027518/1 & MR/W002426/1). A.C.M & J.V.W. are funded by a Wellcome Trust Collaborative Award (224257/Z/21/Z).

### Authors’ contributions

Author contributions: G.G. conceived, designed, and performed experiments, analyzed data, and wrote the manuscript. F.D. provided conceptual input to the manuscript and edited the manuscript. C.H. performed histology and reviewed manuscript. A.C.M and J.W. analyzed histological data and reviewed manuscript.

## Acknowledgements

We thank Dr Jadwiga Zarebska, Dr Bryony Scott and Dr Vicky Batchelor for providing training on the PMX model, weight bearing behavior and histology. We thank Dr Luke Pattison for assistance with the intra articular injection technique.

## Authors’ information

This research was funded in whole or in part by UK Research & Innovation and the Wellcome Trust. For the purpose of Open Access, the author has applied a CC BY public copyright license to any Author Accepted Manuscript (AAM) version arising from this submission.

## Notes

https://osf.io/fg9jx/files/osfstorage/66cf3edada345a296e8c9591

### Competing Interest Statement

The authors have declared no competing interest.

### Summary of Updates

Revised introduction and discussion; Included new statistical analysis of light phase homecage data; Supplementary figure 1 is now figure 2; new supplementary figure

