## Supplementary data for "Using in vivo calcium imaging to examine joint neuron spontaneous activity and home cage analysis to monitor activity changes in mouse models of arthritis"

**Supplementary Table 1.**

| | Primary outcome measure | Statistical test | # Groups | Animals / group | Observed effect size | Sensitivity analysis for row sample size/test |
|---|---|---|---|---|---|---|
| **AIA knee swelling** | Size across knee joint | Independent samples t-test | 2 | 11 | d=2.5* | d=1.26 |
| **PMX histology** | Cartilage loss | RM-ANOVA, between-group effect of treatment | 2 | 9-10 | F=1.04* | F=0.56 |
| **AIA imaging** | % joint silent nociceptors spontaneously active | Kruskal-Wallis test | 2 | 10-11 | F=0.72* | F=0.66 |
| **PMX imaging** | % joint silent nociceptors spontaneously active | Kruskal-Wallis test | 2 | 9-10 | F=0.08 | F=0.66 |
| **AIA behaviour** | Distance travelled<br>Time spent mobile | RM-ANOVA, between-group effect of treatment | 2 | 9-10 | F=1.27* | F=0.55 |
| **PMX home cage behavior** | Distance travelled<br>Time spent mobile | RM-ANOVA, between-group effect of treatment | 2 | 10 | F=0.37 | F=0.52 |
| **PMX Weight bearing** | Hind limb weight bearing ratio | RM-ANOVA, between-group effect of treatment | 2 | 10 | F=0.39 | F=0.50 |

Summary of statistical tests and n-numbers used and the effect sizes we observed. A star indicates when an effect size was statistically significant in a given experiment. The last column provides a sensitivity analysis, indicating the minimum effect sizes that can theoretically be detected with 80% probability when carrying out an experiment with the statistical test and n number provided for each row.

**Supplementary Table 2.** ***In vivo*** **calcium imaging data presented per neuron.**

| | AIA | CTRL | PMX | Sham |
|---|---|---|---|---|
| **# L4 neurons** | 3483 | 4082 | 3197 | 3124 |
| **# Spontaneously active L4 neurons** | 414 | 379 | 227 | 284 |
| **# FB+ neurons** | 63 | 61 | 75 | 55 |
| **# Spontaneously active FB+ neurons** | 14 | 1 | 2 | 2 |
| **Proportion of spontaneously active L4 neurons still active post-lidocaine** | 62/3179 | 64/2522 | 51/3197 | 36/2430 |
| **Proportion of spontaneously active FB+ neurons still active post-lidocaine** | 1/11 | 0/1 | 0/2 | 0/2 |

FB = fast blue. AIA = Antigen induced arthritis. PMX = partial medial meniscectomy

**Supplementary Table 3. Responses of spontaneously active fast blue joint neurons to capsaicin**

| | Spontaneously active prior to capsaicin application | Blocked after capsaicin | Activated and blocked after capsaicin | Activated by capsaicin |
|---|---|---|---|---|
| **# neurons** | 8 | 6 | 2 | 2 |
| **%** | | 75.0% | 25.0% | 25.0% |

Data from n=5 mice that had capsaicin applied to the nerve.

**Supplementary data**

Supplementary Video 1.avi
https://osf.io/fg9jx/files/osfstorage/66cf3edada345a296e8c9591
**Supplementary Video 1. Example recording of a spontaneously active fast blue labelled joint neuron in the AIA model.** The bottom panel shows an area taken from the maximum intensity projection of a z-stack. The top panel shows a timelapse recording of GCaMP6s activity in the corresponding area. The fast blue labelled joint neuron with spontaneous activity is indicated by the arrows. Time-lapse recorded at 3.65Hz. Scale bar = 50 µm

**Supplementary Figures**

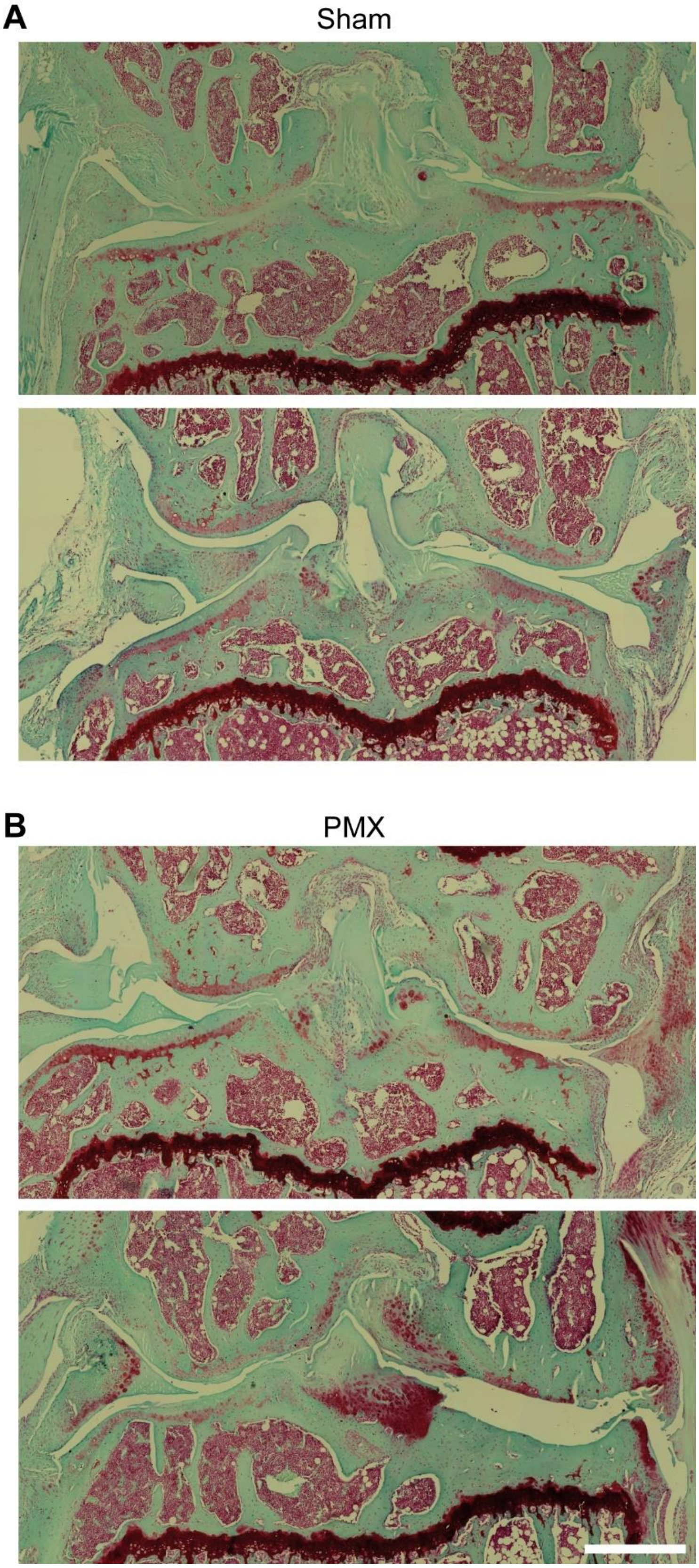


**Supplementary Figure 1. Additional examples of joint histopathology in sham and PMX mice.** Example images of Safranin O & Fast Green-stained knee joint sections in Sham (A) and PMX (B) mice at 13-15 weeks post-surgery. Scale bar = 500µm.

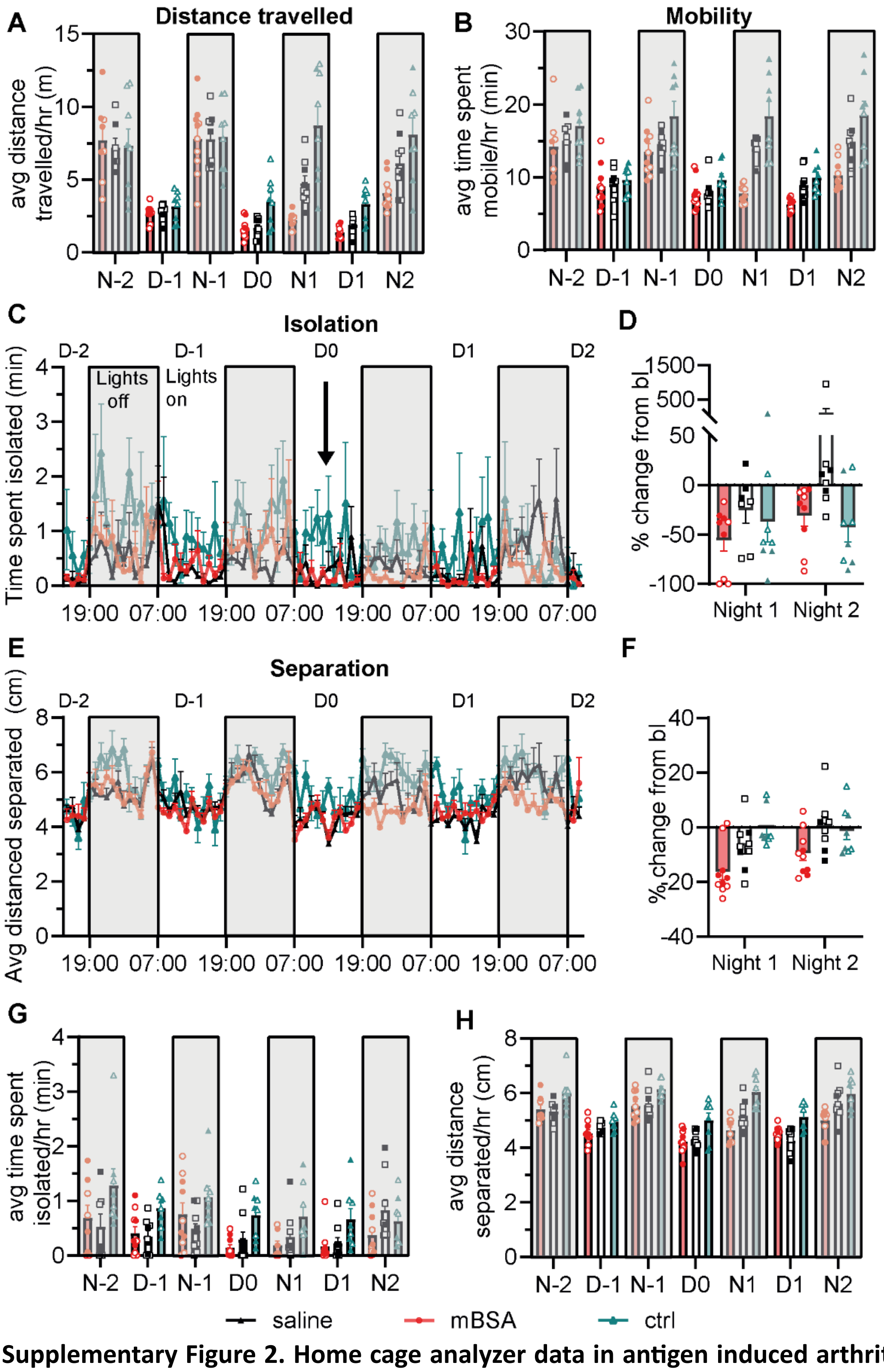


**Supplementary Figure 2. Home cage analyzer data in antigen induced arthritis mice.** A&B) Graphs showing distance (A) and mobility (B) data presented in 12-hr time bins. C-E) Graphs showing the time spent isolated (C&D) and the average distance separated (E&F) before and after intraarticular injections (indicated by arrow in C). Data in C) and E) are presented as mean +/- SEM in 1hr intervals. Grey bars represent the dark phase. Data in D) and F) show the percentage change from baseline for the dark phase (19:00-07:00). G&H) Graphs showing isolation (G) and separation (H) data presented in 12-hr time bins. mBSA: n = 10, saline: n = 9. Males = closed circles, females = open circles.

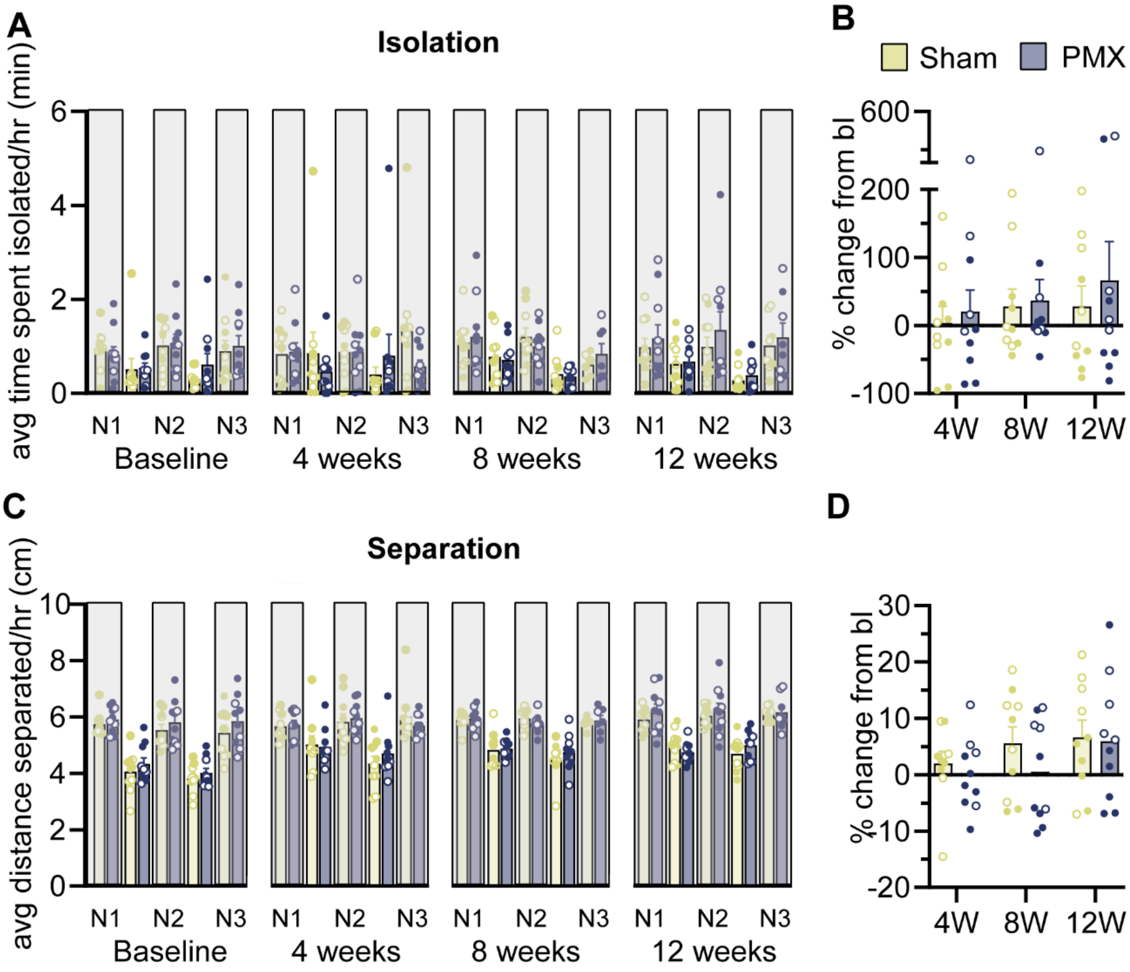


**Supplementary Figure 3. Home cage analyzer data in osteoarthritis mice.** A-D) Graphs showing the time spent isolated (A&B) and the average distance separated (C&D) at baseline and at various time intervals post-PMX surgery. Data in A) and C) are presented as mean +/- SEM in 12hr intervals. Grey bars represent the dark phase. Data in B) and D) show the percentage change from baseline for the dark phase (19:00-07:00). PMX: n = 10, sham: n = 10. Males = closed circles, females = open circles.

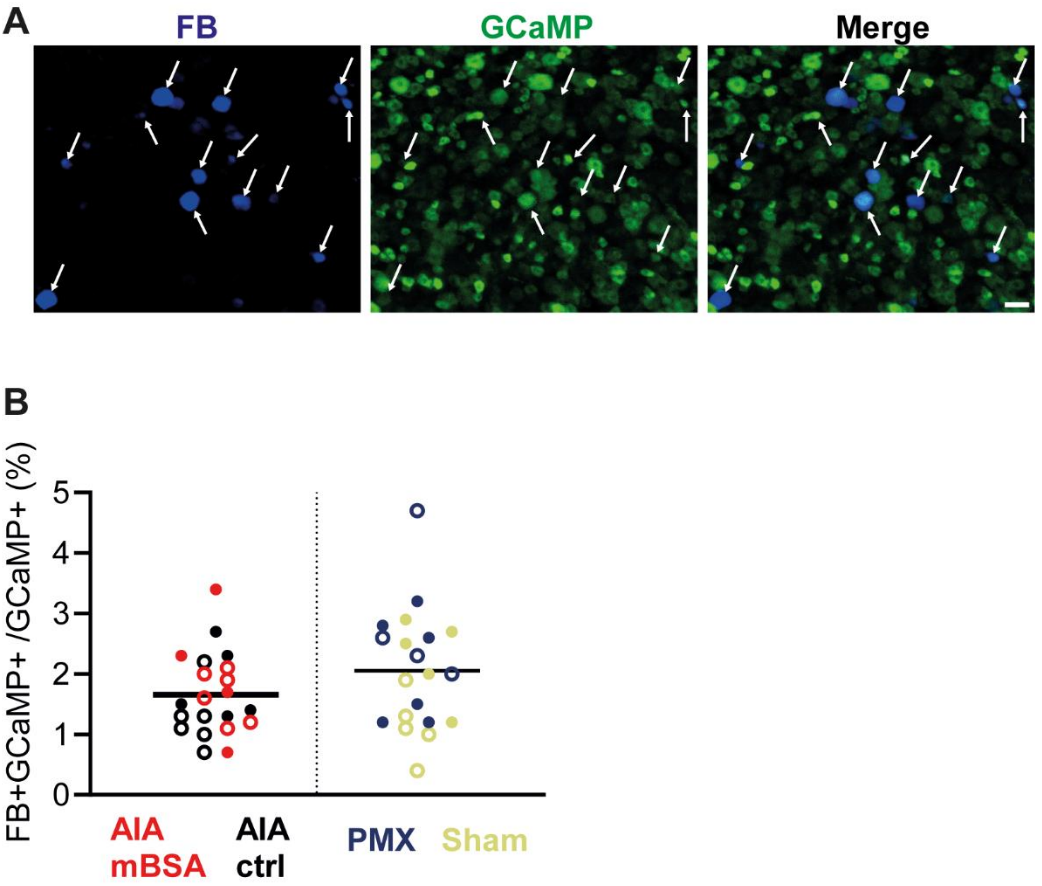


**Supplementary Figure 4. The proportion of GCaMP6s+ neurons labelled with fast blue was similar in AIA and PMX mice.** A) Example image showing max projection from a z-stack taken in blue (fast blue) and green (GCaMP6s) channels on the confocal microscope. Arrows indicate neurons that are labelled with fast blue (FB) and GCAMP6s. Scale bar = 50µm. B) Graph showing the proportion of GCaMP+ neurons labelled with fast blue for each experimental group. males = closed circles, females = open circles.

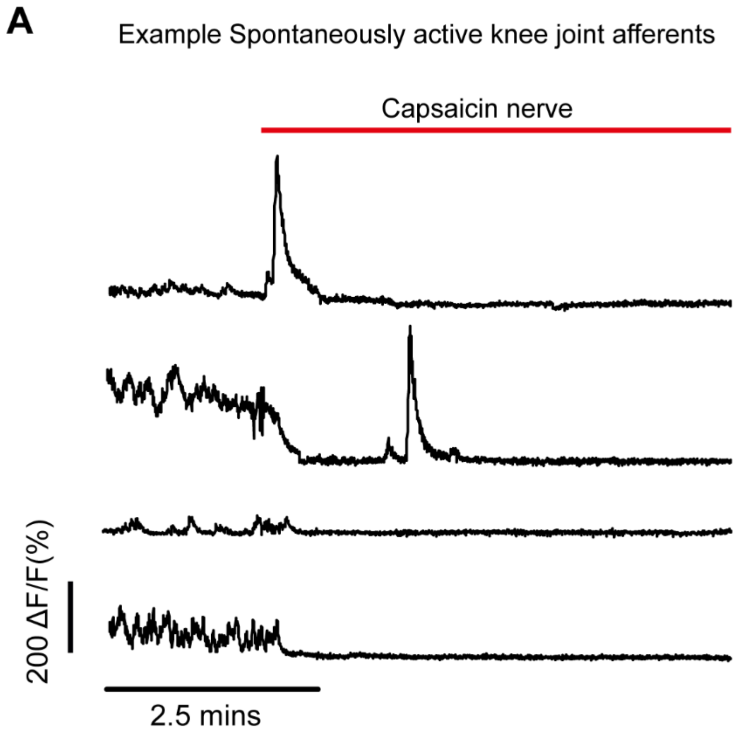


**Supplementary Figure 5. Capsaicin application to the nerve produced variable responses.** A) Example traces from spontaneously active joint neurons before and after application to the nerve. Note that activity in some neurons was blocked following capsaicin-induced depolarization (upper trace) whereas others were blocked without any prior depolarization (lower two traces).

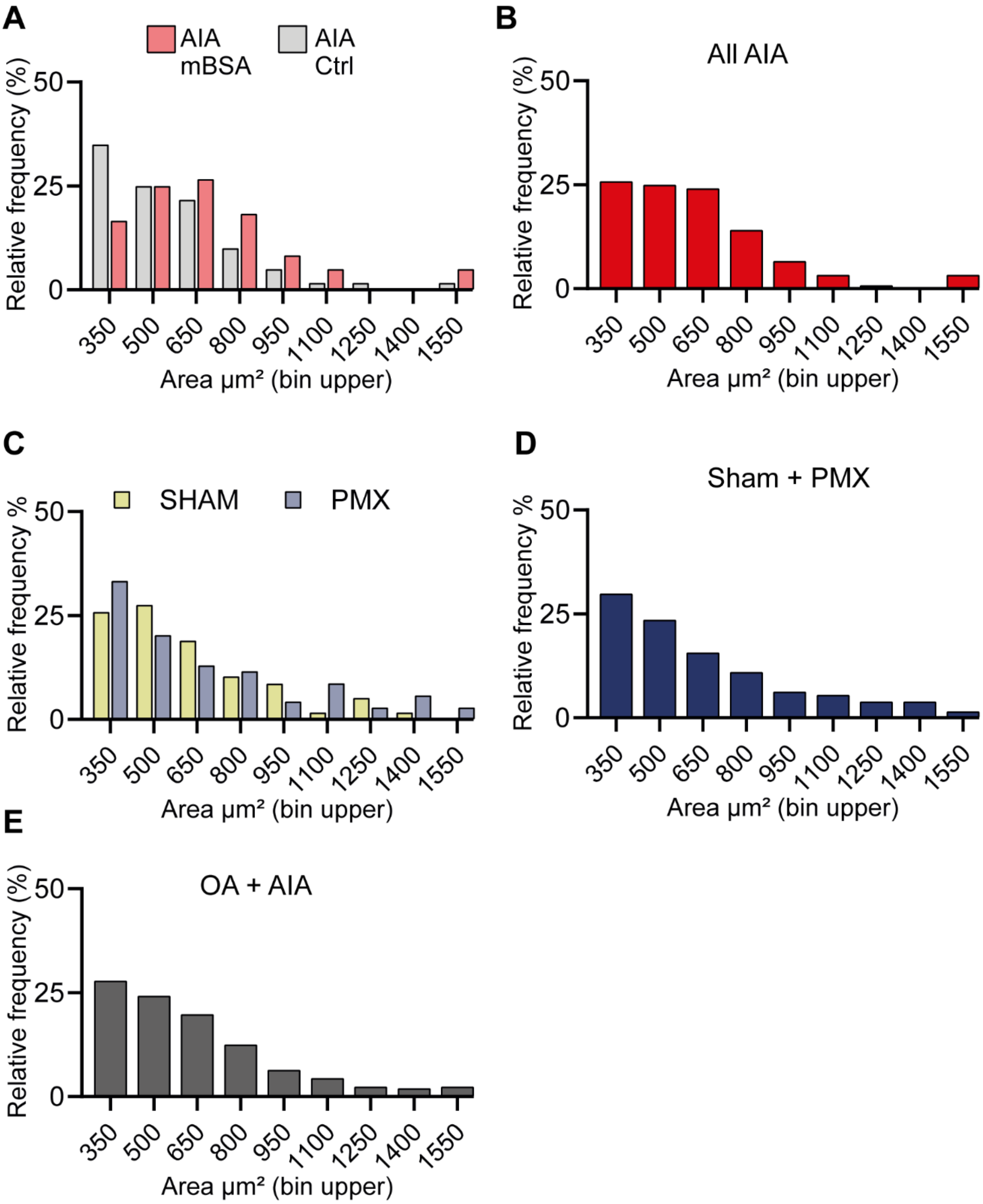


**Supplementary Figure 6. Size histograms for antigen induced arthritis and partial medial meniscectomy mouse cohorts.** A-E) Size histogram showing the relative frequency of fast blue (FB+) joint neurons in bin widths of 150 µm². AIA cohorts: AIA mBSA = 63 neurons from n = 10 mice. AIA Ctrl = 61 neurons from n = 11 mice. All AIA = 124 neurons from n = 21 mice. PMX cohorts: Sham = 58 neurons from n = 10 mice. PMX = 71 neurons from n = 10 mice. SHAM+PMX = 129 neurons from n = 20 mice.
