## Supplementary methods for "Using in vivo calcium imaging to examine joint neuron spontaneous activity and home cage analysis to monitor activity changes in mouse models of arthritis"

**Data loss and exclusion**

We aimed to collect data from n=12 mice/group for AIA and PMX cohorts. However, some data was lost or excluded for reasons outlined below.

Data losses: One mouse from the AIA cohort had to be culled for a welfare reason. Two mice from the PMX cohort were culled due to welfare reasons. Two mice from one batch of OA model mice had ear marks that could no longer be differentiated at the end of the study. As these mice were from different treatment groups, all the data collected from these mice was disregarded.

Data exclusion: Initially, AIA experiments were performed by injecting volumes of 10µl into the knee joint. Following injections with these volumes, we observed abnormal sensitization i.e. spontaneous activity of many sensory neurons in both saline and mBSA groups. After the experimenter responsible for the injections received further training from an expert, Dr Luke Pattison at Cambridge University, abnormal neuronal activity was rarely observed. We speculate that the tibial nerve, which sits at the back of the knee may have been damaged by holding the knee too firmly during these initial injections. More care was taken thereafter, and saline and mBSA intra-articular injection volumes were reduced to 2.5µl in an abundance of caution to avoid damaging the small joint capsule space with high volumes of liquid. In these initial experiments, we also observed that following injections of 5µl fast blue, the muscle surrounding the knee joint was stained yellow suggesting that the fast blue had leaked out of the joint. Therefore, in future experiments we only injected a maximum of 2µL of fast blue into the joint, which aligns with the reported maximum volume you can inject into the joint without it leaking from the joint capsule (48). Consequently, only data collected with the new injection volume regime (2.5µl mBSA, 2µL fast blue) were included in the analyses presented here.

For knee joint histology experiments, one sham sample was not included within the analysis due to an embedding error that caused an issue with orientation that could not be resolved by re-embedding.

For home cage experiments, data were excluded from one batch of male AIA mice because of the loss of one mouse prior to the experiment starting (see above) and a faulty microchip in another mouse. Thus, since data from only n=2 out 4 mice was collected for this batch, the home cage data (and environment) was unlike other batches, so it was not analyzed.

For imaging experiments, 1 AIA mBSA mouse was excluded due to poor GCaMP6s labelling and 1 sham mouse in the OA cohort was excluded from the fast blue analysis because only 1 cell was labelled.

In some imaging experiments (n=3 for OA cohort and n=5-6 for AIA cohort, a spring-loaded clip was used to apply mechanical stimulation (~450g) across the knee joint. We used a high force in attempt to reveal unmasking of silent nociceptors in the RA and OA models. However, the results from this relatively small set of mice were variable and showed that not all joint afferents are activated by applying mechanical stimulation in this way; we therefore did not include these data in this manuscript. Interested parties can download the traces from the SPARC portal if they wish. We are currently optimizing different methods of activation of mechanically-sensitive joint afferents I.e. flexion/extension and torsion and intend to publish the clip data with other related data in a future manuscript.
